# Inflation versus filling-in: why we feel we see more than we actually do in peripheral vision

**DOI:** 10.1101/263244

**Authors:** Brian Odegaard, Min Yu Chang, Hakwan Lau, Sing-Hang Cheung

**Author notes:** Authors for correspondence: (1) Brian Odegaard, (2) Sing-Hang Cheung. Present address (1) Franz Hall, 502 Portola Plaza, Los Angeles, CA 90095 (2) 6th Floor, Jockey Club Tower, The University of Hong Kong, Hong Kong.

## Abstract

Do we perceive fine details in the visual periphery? Here, we propose that phenomenology in the visual periphery can be characterized by an *inflated* sense of perceptual capacity, as observers overestimate the quality of their perceptual inputs. Distinct from the well-known perceptual phenomenon of “filling-in” where perceptual content is generated or completed endogenously, inflation can be characterized by incorrect introspection at the subjective level. The perceptual content itself may be absent or weak (i.e., not necessarily filled-in), and yet such content is mistakenly regarded by the system as rich. Behaviorally, this can be reflected by *metacognitive* deficits in the degree to which confidence judgments track task accuracy, and *decisional* biases for observers to think particular items are present, even when they are not. In two experiments using paradigms which exploit unique attributes of peripheral vision (crowding and summary statistics), we provide evidence that both types of deficits are present in peripheral vision, as observers’ reports are marked by overconfidence in discrimination judgments and high numbers of false alarms in detection judgments. We discuss potential mechanisms which may be the cause of inflation and propose future experiments to further explore this unique sensory phenomenon.

## Introduction

How much of the visual periphery do we actually see? Some findings indicate that we perceive the periphery in precise detail [1] and that limitations in our ability to recall items are based mainly on memory, rather than sensory, processing constraints [2,3] But findings such as inattentional blindness [4] and change blindness [5,6] indicate that perception of items presented outside the fovea is quite limited. Thus, a question arises as to whether our subjective sense of the visual periphery is *inflated* beyond what we should expect based on the underlying processing limitations.

Research on two visual phenomena present unique opportunities to explain the puzzle of peripheral phenomenology: crowding and summary computations. Crowding is defined by deficits in the ability to identify objects surrounded by “clutter” in the visual surround [7]. For example, identifying the middle letter in a row of three letters is relatively easy when they are presented in the center of the visual field, but surprisingly difficult when they are shown in the periphery. Results reveal that crowding can change appearance [8,9], and therefore may be at least partially responsible for impairments in identifying objects in the periphery. Crowding can even result in metacognitive errors [10], indicating that it affects not only perceptual performance, but likely subjective phenomenology as well.

Summary statistics are defined by the visual system’s tendency to represent fine details in the visual periphery as an ensemble, as individual components are compressed into a gist-based representation [11]. This capacity for summary representation extends across a wide variety of dimensions, as observers can estimate average size [12], motion direction [13], position [14], and orientation [15] of groups of elements quite effectively. It has been posited that summary statistics may underlie phenomenological experience of the visual periphery [16]. This view finds support in work using metamers [17], which shows that pooling mechanisms outside the fovea can cause distinct images to be perceptually indistinguishable. This demonstrates how distortions of peripheral visual content may not always result in subjective perceptual differences. And yet, introspectively, we don’t seem to think we would fail to notice such distortions.

How can we characterize this mismatch between introspective phenomenology and actual perceptual representational quality in peripheral vision, to go beyond anecdotal descriptions? Traditionally, the mechanism of “filling-in” is thought to be important and relevant. Filling-in is a perceptual phenomenon whereby features from surrounding regions of the visual field are perceived despite their physical absence in a particular location [18]. Typically, this is thought to be achieved by having the perceptual content in early sensory systems (e.g V1) generated endogenously [19]. That is, actual content is created in the absence of external input. This can lead to illusory perception of color [20], texture [21], motion [22], brightness [23], and other visual attributes. Filling-in is most evident in the blind spot, where the visual system compensates by representing similar content in this region without inputs [24], but is also evident in perceptual illusions like neon color spreading [25] and the Troxler effect [26]. Evidence indicates that the neural mechanisms underlying filling-in reside in early-level visual areas [18,27–29], as early sensory representations are completed based on top-down rather than bottom-up input.

However, over and above the degree to which filling-in may play a role across the visual field, we hypothesize that a second process, *inflation*, also plays a role in perception of the visual surround. Inflation can be defined as the subjective overestimation of the reliability or quality of the sensory representations themselves. That is, the representations themselves are not necessarily filled-in with details, but are subjectively misestimated to be rich in content. Across the entire visual periphery, it is unlikely filling-in processes provide all the fine details in early sensory regions in a precise, pixelated representation instantly, as soon as we view a scene. In addition, there is evidence that even in cases where filling-in occurred, such as in the blind spot, there are additional subjective biases to be accounted for [30].

To assess inflation in experiments, there are two aspects of behavior that can be investigated: one may show a *metacognitive* bias to be overly confident in perceptual judgments [31] given current processing limitations, and/or a *decisional* bias to think things are present, even when they are not [32]. Because these biases can be captured with signal detection theoretical (SDT) measures, they can be readily characterized in quantitative terms in psychophysical experiments. Importantly, just because these biases are in terms of decision or confidence criteria does not mean they only reflect shifts in response strategy; it has been argued that these biases can reflect subjective perceptual phenomenology, which we interpret is likely the case here as well [33,34]. In part, this argument is due to the observation that feedback and training did not seem to remove such biases [31]; if they were at the cognitive or response level, we would expect them to be more flexible and adaptive.

Previous work has already provided support for this inflation account for stimuli perceived under lack of attention. For example, according to [31], under conditions of inattention, representational precision of visual information is reduced, but a similar criterion is used compared to attended conditions, resulting in higher numbers of false alarms when making detection judgments [32], and higher ratings of visibility when making discrimination judgments [31]. Inflation can be interpreted to follow similar principles. In the visual periphery, processing capacity is reduced [35,36]. Similar to what has been shown under inattention, this may lead to an overestimated sense of how visible the periphery is, despite deficits in processing.

Here, to investigate the role that inflation may play in the periphery, we combined the study of crowding and summary statistics with SDT to quantitatively characterize whether inflation occurs in each of these scenarios. Specifically, we will show how the methods of assessing metacognitive bias (in a discrimination task) and detection biases may be useful in different contexts. In our crowding study, we assessed whether metacognition in a discrimination task is impaired in the periphery. This is because crowding typically impairs discrimination but not detection itself [37]. We aimed to evaluate confidence judgments in crowded conditions to investigate how effectively confidence tracks performance on a trial-by-trial basis. On the other hand, in our summary statistical study, we evaluated whether there may be detection biases when subjects had to detect line patches with a coherent orientation pattern. One advantage of detection tasks is that we can look at the false alarm trials, where the physical stimuli are matched between the central and the peripheral locations. To compare performance between center and periphery, one often ends up using stronger stimuli for the periphery, for otherwise sensitivity may be too low compared to central vision. In focusing only on false alarms, we bypass this issue of stimulus confound.

## Experiment 1: Metacognition in Crowding

In our first experiment, we explored how crowding, an omnipresent phenomenon in the visual periphery in everyday settings, may be linked to inflation. Specifically, we were interested in whether trial-by-trial confidence ratings would effectively track task performance, or whether these ratings would reveal impaired metacognition for elements in the visual surround (see Figure 1).

**Figure 1.**
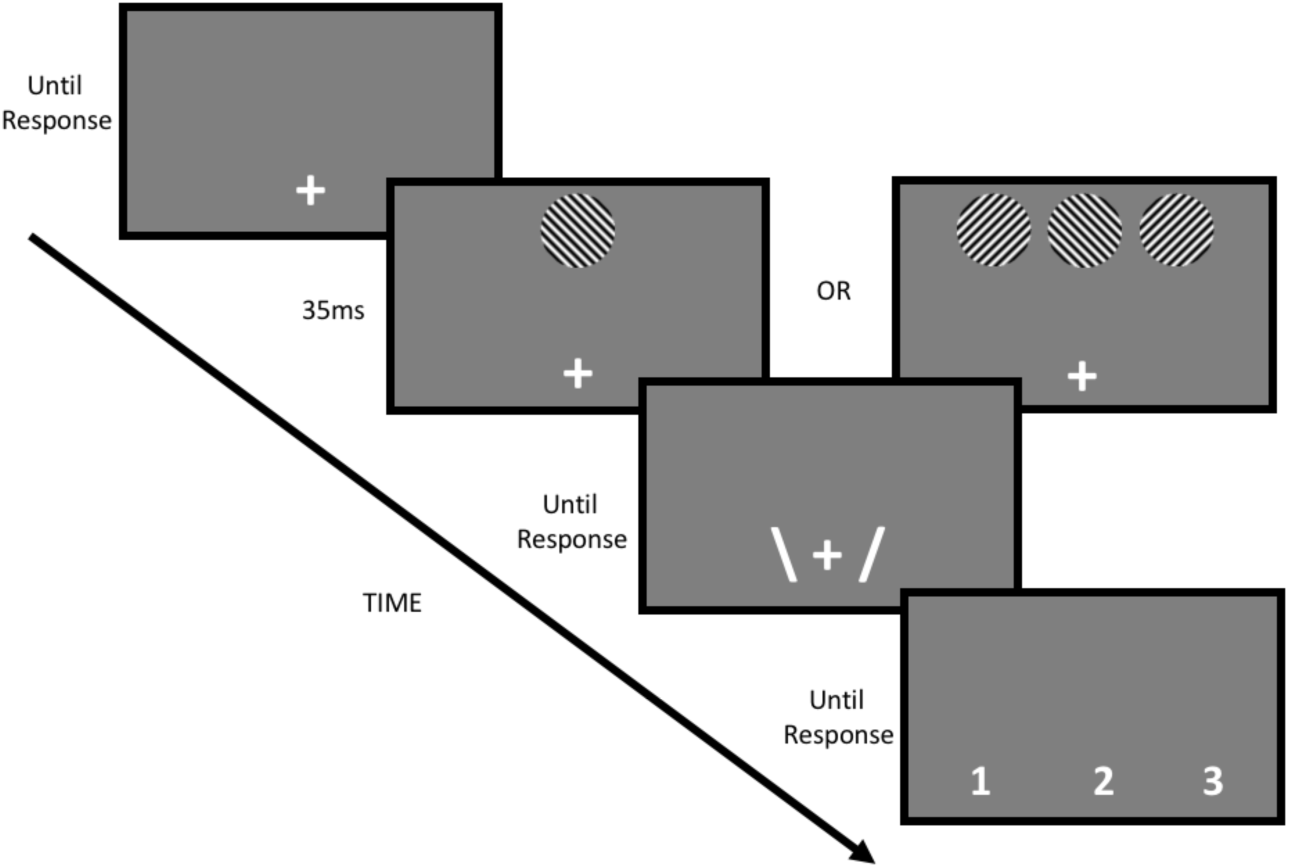
Protocol for each trial in Experiment 1. The fixation cross was shown throughout the trial. After participants pressed the ‘space bar’ on the keyboard to initial the trial, the target sine wave gratings were presented above the fixation cross. This target grating could either be presented alone (single condition), or surrounded by other gratings on each side (crowded condition), and all patches were presented for 35ms. Participants then had to report the orientation of the target patch (left or right) and also rate their confidence for their report on a scale of 1-3. We note that the sine wave gratings displayed in this figure are not to scale; we increased their size to improve appearance, but see Methods for details about size.

## EMethods

### Participants

Thirty young adults (18-30 years old, M = 22.00, SD = 2.95; 25 females) with normal or corrected-to-normal vision were recruited from the University of Hong Kong to participate in this experiment. All participants volunteered and received no monetary compensation for their time spent in the experiment. This experiment was part of the second author’s undergraduate thesis study and was approved by the Departmental Research Ethics Committee in the Department of Psychology at the University of Hong Kong. Informed consent was obtained from all participants before the experiment began. Twenty-three participants successfully completed this task. Among the seven participants that were excluded, five were excluded due to very low threshold differences between the crowded and single conditions, which indicates absence of crowding (possibly due to unstable fixations), and two were excluded due to not following the instructions (one exhibited near-chance accuracy [< 60%] and one exhibited a negative meta-d’ score).

### Apparatus and Materials

Participants attended the experimental session in the Department of Psychology at the University of Hong Kong. The experiment was coded in MATLAB using the Psychophysics Toolbox [38–40] and custom-written code for stimulus presentation. Stimuli were presented on a 17-inch CRT monitor (1024 × 768 pixel resolution at 85Hz refresh rate). Background luminance was 17.8 cd/m^2^ with ambient light turned off. A headrest and a chinrest were used to help the participants maintain a viewing distance of 92cm.

Both target and flankers were sine wave gratings (2 cpd) presented through a circular window of 2.5° in diameter. Orientation of the gratings was either 45° clockwise or 45° counterclockwise; both orientations had an equal probability of being displayed on a given trial. The target was presented 10° above the fixation cross, which subtended 0.3° and was presented near the bottom of the display. Two flanker gratings were presented left and right of the target at a target-flanker distance (center-to-center) of 3° in the crowded condition. This combination of target eccentricity and target-flanker distance was based on previous paradigms which showed robust crowding [41,42].

### Procedure

After signing the informed consent form, participants were introduced to the task (Figure 1). Participants performed an orientation discrimination task in each trial. Each trial started with the participants fixating on the fixation cross. Participants pressed (Space) to initiate stimulus presentation. Either the target alone (single condition) or the target with two flankers (crowded condition) were then presented for 35ms. After the stimulus screen, two vertical lines tilted clockwise and counterclockwise were presented to the right and left of the fixation cross, respectively, to prompt the participants to respond using the number pad. Participants pressed (4) for left or (6) for right, and no feedback was given. After the orientation judgment, participants also reported their confidence in their judgment using a scale from 1 (not at all confident) to 3 (extremely confident). Participants pressed (4) for 1, (5) for 2 and (6) for 3 in rating their confidence. We did not monitor eye movements in this experiment. However, the exposure duration we used was too short to execute a saccade from the fixation cross to the target.

Each trial block consisted of 64 single and 64 crowded trials. We used separate fixed-step-size staircases to continuously adjust the Michelson contrast levels for the two types of trials. A one-up one-down staircase with a down-step size to up-step size ratio of.2845 was chosen to achieve a target accuracy level of 77.85% [43]. There were two blocks of practice trials, followed by eight blocks of experimental trials. Participants performed only the orientation discrimination task during the first practice block. Initial contrast levels were set at .4 and .8 for the single and crowded trials respectively in the first block. Staircases in blocks two to ten started with final contrast levels from the previous block. The signal detection theoretic measures *d’* and meta-*d’* [44] were calculated based on blocks three to ten. If the staircases worked as planned, *d’* should be matched between the single and crowded conditions. However, our current setup failed to render the required contrast levels (i.e., contrast was not low enough for the single condition or required contrast going beyond 1 for the crowded condition) for some participants. Therefore, we observed a statistically significant difference in *d’* between the single and the crowded conditions.

### Analysis

Each participant’s data were analyzed using custom software for Signal Detection Theory (SDT) analysis [44–46]. Specifically, we used the fit**_**meta**_**d**_**MLE.m file to estimate both Type 1 (*d’*) and Type 2 (meta-*d’*) SDT parameters for sensitivity.

## Results

As shown in Figure 2, participants displayed better performance (measure by the signal detection theoretic measure *d’*) in the single condition compared to the crowded condition, t(22) = 7.35, p < 10^−6^. Interestingly, participants displayed relative metacognitive impairments in the crowded condition compared to the single condition. We used the M-ratio to quantify metacognitive efficiency. The M-ratio, which is the fraction meta-*d’*/*d’*, represents the amount of signal strength available for metacognition, and reflects the metacognitive efficiency in a given subject [44,46–48]. An M-ratio near 1 represents metacognitively ideal performance. As can be seen in Figure 2, on average, participants were close to metacognitively optimal in the single condition; the small exceedance above 1 can be ascribed to estimation error or that they did not perform the primary discrimination task perfectly according to SDT. However, in the crowded condition, participants displayed clear metacognitive deficits, as the M-ratio was significantly lower in this condition compared to the single condition, t(22) = 4.26, p < 10^−3^. Thus, when experiencing crowding effects in the visual periphery, subjective assessments of how we see deviate from optimality. One could argue this may be due to the fact that in the crowded conditions, *d’* itself was lower, and the M-ratio method may not have removed the influence of this difference perfectly, but the next result addresses this concern.

**Figure 2.**
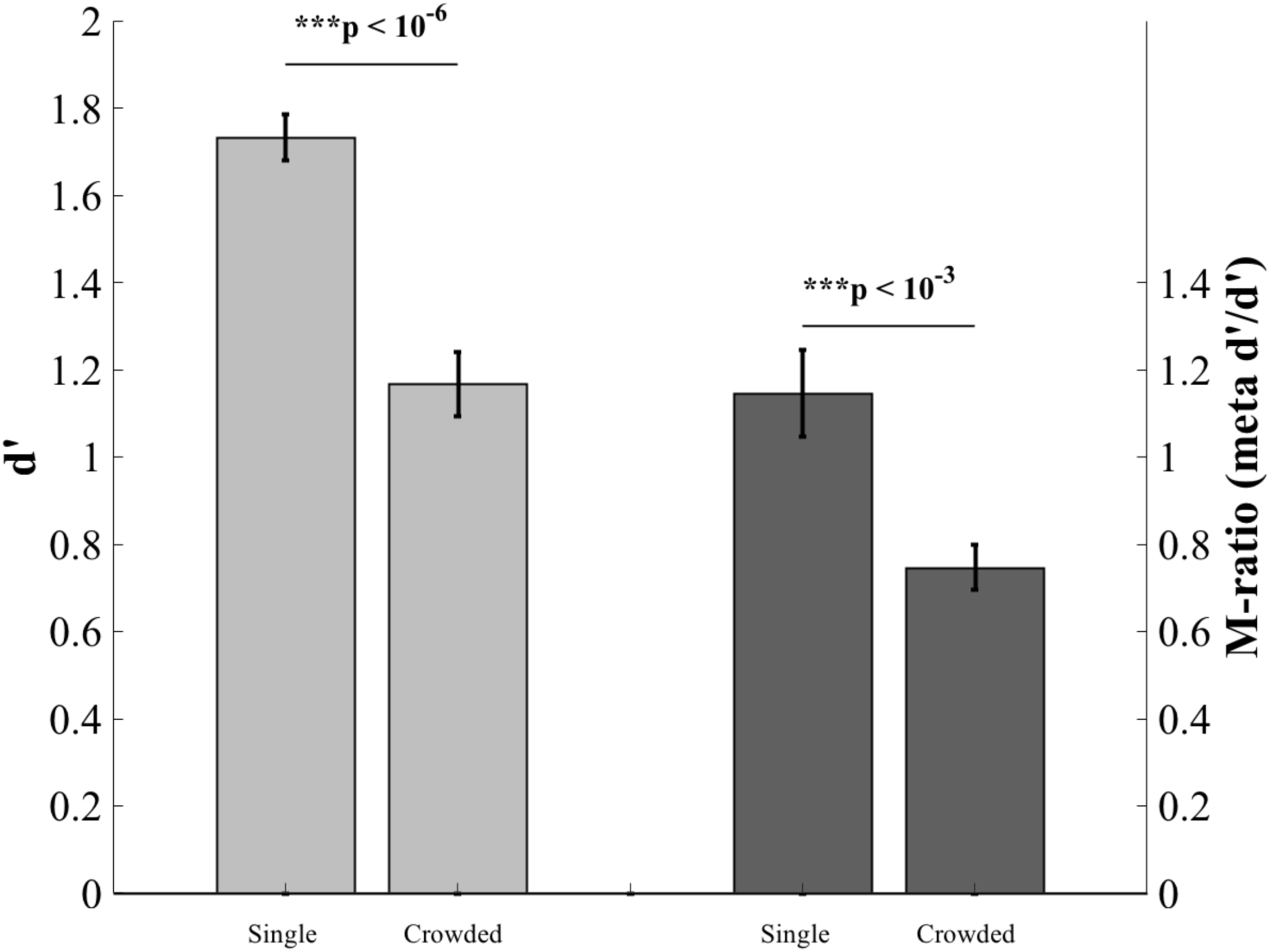
Perceptual sensitivity and metacognitive efficiency in an orientation discrimination task. Shown here are the results from 23 participants in Experiment 1. As shown by the light gray bars, participants were much less effective at discriminating the orientation of a tilted grating compared when it was surrounded by other gratings, compared to discriminating the orientation of a single grating. *d’* is the standard detection-theoretic measure of sensitivity. Shown by the dark grey bars is a measure of the metacognitive efficiency (the M-ratio; meta-*d’* / *d’*) in both conditions, which indicates how effectively confidence ratings could distinguish between correct and incorrect judgments. As can be seen in the figure, metacognitive efficiency was impaired in the crowded condition compared to the single condition.

To better illustrate the basis of this phenomenon, we also analyzed average confidence in the single and crowded conditions, separating trials by whether they were correct or incorrect (Fig. 3). As can be seen in this figure, on *correct* trials, confidence was approximately the same between the single and crowded conditions, t(22) = -0.25, p = 0.81. However, on *incorrect* trials, confidence was higher for the crowded trials compared to the single trials, t(22) = -8.46, p < 10^−7^. Notably, this higher confidence was shown despite the fact that people were overall less accurate in the crowded condition. Therefore, the deficit in metacognition in crowding seems to be primarily driven by overconfidence on incorrect trials: when participants are wrong about what they see in the periphery, they don’t always know it, and have more confidence in their perceptions than what is warranted.

**Figure 3.**
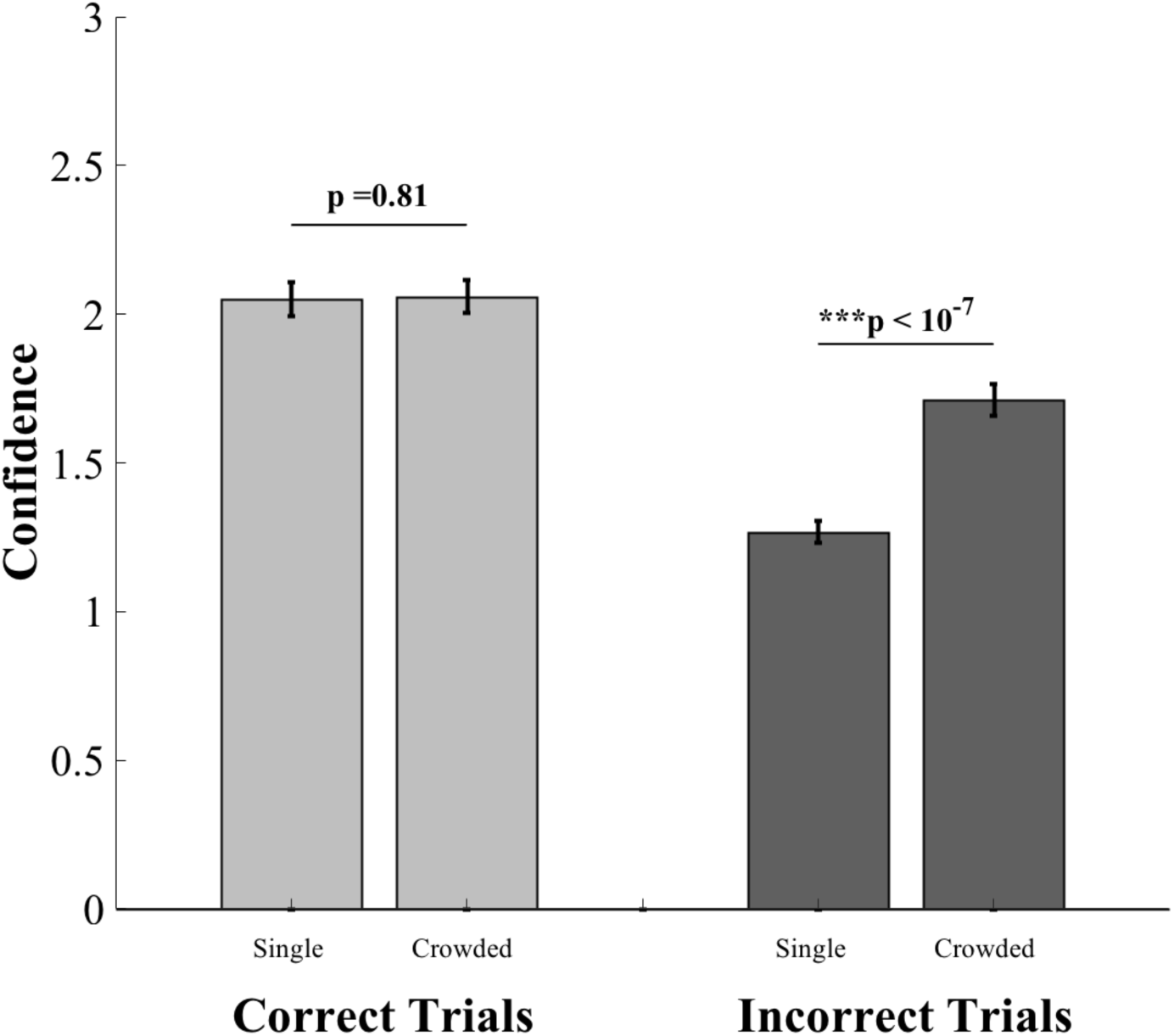
Average confidence for correct and incorrect trials. The light gray bars indicate the average confidence for correct trials for the single and crowded conditions. As can be seen in the figure, the difference between these conditions is not significant. The dark gray bars indicate the average confidence for incorrect trials. A clear difference between the single and crowded conditions is evident, and participants are significantly more confident in the incorrect crowded trials compared to the incorrect single trials.

## Experiment 2: Detection based on summary statistics

It has been proposed that part of what characterizes phenomenological experience of the visual periphery is the visual system’s capacity to represent groups of items as ensembles or gist-based representations [16]. In other words, rather than encoding details in the periphery with high fidelity, visual information is compressed to eliminate redundancy and represent information outside the fovea in the form of summary statistics [11]. Considering the results from Experiment 1, it is an intriguing question how summary statistical judgment may be biased by the fact that individual crowded items may nonetheless provide a subjectively reliable percept, so that all the items together may subjectively look as if they are more coherent than they really are.

In Experiment 2, we investigated whether such detection biases exist in the periphery. On each trial, we presented observers with a diamond-shaped stimulus composed of many individual lines with various orientations (Fig. 4). We hypothesized that, similar to previous investigations, observers would be much more likely to say that a congruent patch of lines was *present*, even when the lines were composed of only random orientations.

**Figure 4.**
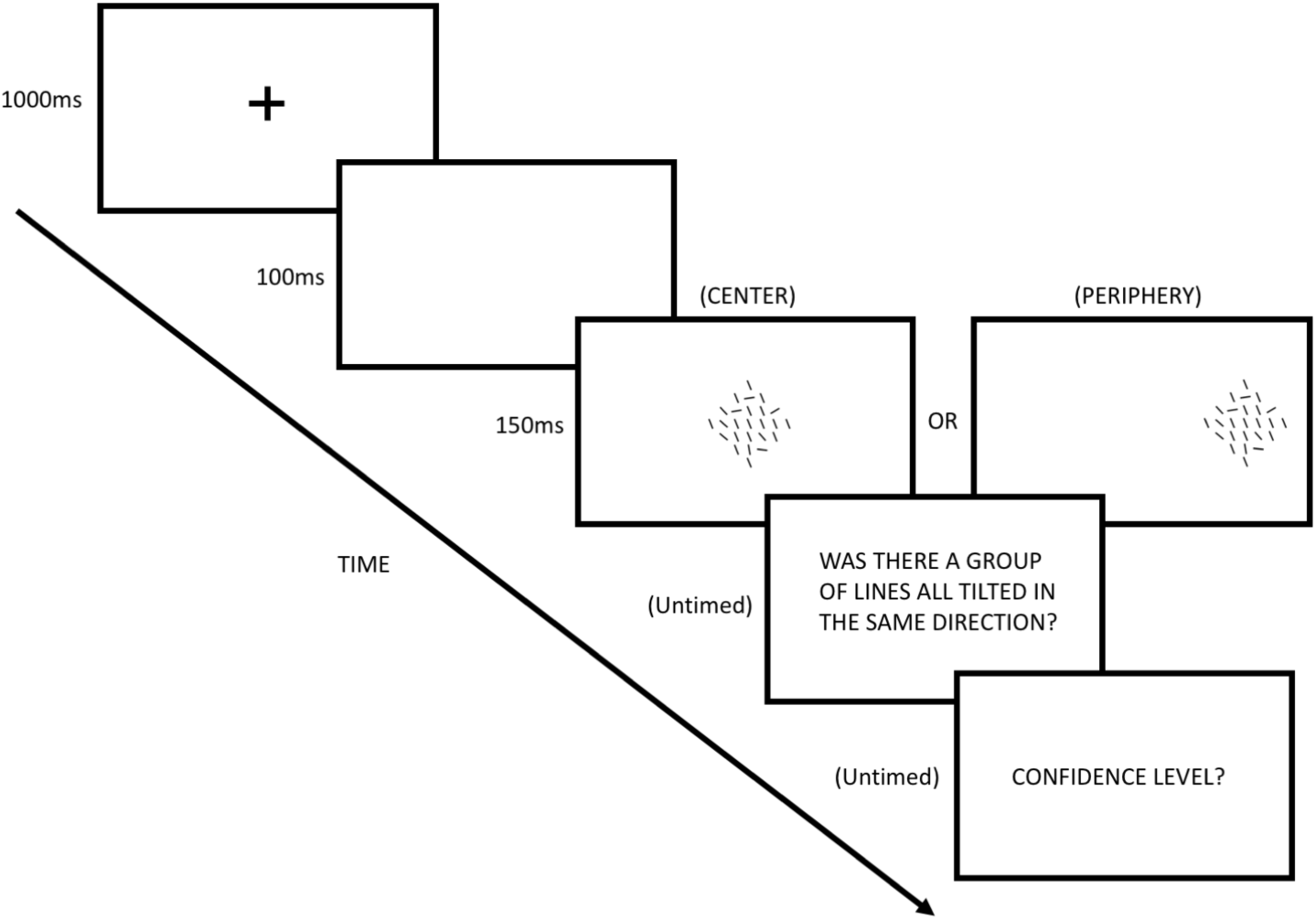
The protocol for each trial in Experiment 2. Each trial began with a fixation cross for 1000ms, followed by a 100ms blank screen. Then, lines were presented at either a central or peripheral location for 150ms. Following presentation of the lines, participants responded whether a group of coherent lines with the same orientation was present, and gave their confidence on a scale of 1-4. In this example, there are 16 lines with congruent orientation in the image. Please note that the wording shown in this schematic differs slightly from the actual wording displayed in the experiment.

## Methods

### Participants

Seventy-four total research participants responded to an advertisement on Amazon’s mTurk online platform and successfully completed the experimental task. Three participants were excluded due to errors in the fitting procedure from the MATLAB files for estimating signal detection parameters [45]. No personal or demographic information about participants was collected, with the exception of using each participant’s unique Amazon mTurk ID to process payments. Research participants were informed before the study that it would require approximately one hour to complete, and that they would earn $4 upon finishing the task, with the possibility of earning an additional $1 bonus if their performance on the task was better than the previous participant. Participants were notified that they could drop out of the experiment at any time, and were informed that they would be paid at a prorated amount of $1 per 15 minutes for the amount of time they participated in the study.

### Apparatus and Materials

We required all participants to use Google Chrome as their web browser for the experiment by adding code which excluded other browsers from running the task. Participants were informed of this requirement before beginning the experiment. The experiment was coded in JavaScript using plugins from the jsPsych library [49] and custom-written code for stimulus presentation. The psiTurk platform [50] was used to launch the study, administer subject payments, and control various elements of the task presentation and design (e.g the hours when the task could be completed, the maximum time allowed to complete the task, enforce U.S. IP addresses for participants, and other details).

### Procedure

Following acceptance of our online “HIT” (Human Intelligence Task) advertisement on Amazon’s mTurk website, participants were presented with a consent form for the experiment, which was approved by the UCLA Institutional Review Board (#15-001484). Once participants agreed to the terms in the consent form, a new browser window was opened and participants began the main experiment. First, instruction screens were presented to request that participants be seated approximately one arm’s length away from their computer screen, and to be positioned directly in front of the screen. Next, participants were informed of the experimental task.

Participants were instructed that they would be required to make judgments about a diamond-shaped pattern of twenty-five black lines drawn on a white background. Each line was 4 pixels wide and 30 pixels high, and spacing between each line was 37.5 pixels on average, with a small amount of random jitter added to each position. Participants were asked to judge whether there was a group of lines that were all tilted in the same direction, or whether the lines were drawn only with random orientations. On trials where a group of lines with congruent rotations were shown, lines with random orientations were resampled if the randomly-selected orientation was within 10 degrees of the congruent orientation direction. Participants were informed that the group of lines with a common orientation could be any number of lines, and that the lines did not have to be next to one another to be considered part of the group.

The experiment began with practice trials to familiarize participants with the stimuli and task. To begin, three easy practice trials were presented where participants were shown the line stimulus for 2000ms and then asked to indicate whether a group of lines were all tilted in the same direction. Participants pressed (Q) for Yes, and (P) for no, and were given feedback about whether the response was correct.

These three trials were followed by two “practice” blocks of 60 trials each, where staircase procedures with fixed step sizes were implemented. The goal was to establish how many lines with coherent orientations should be presented for easy, medium, and hard levels of difficulty, with these three levels designed to approximate ceiling-level performance, ˜85% correct, and ~71-77% correct, respectively [43,51,52]. In the first practice block, a two-up one-down staircase procedure [51] was implemented to estimate the “hard” level of difficulty. Each trial began with a fixation cross presented at the center of the screen for 1000ms. Following a 100ms blank screen, one group of lines was flashed at the location of the fixation cross for 150ms. 800ms later, another group of lines was flashed for 150ms at the same location. Participants were informed in advance that only one of the two sets of lines contained a group tilted in the same direction, and had to indicate whether the first or second presentation contained the coherent group by pressing (Q) or (P) to indicate the first or second presentation, respectively. Participants were also informed that these practice blocks counted towards whether they earned the bonus, to increase incentive to put forth effort as the staircase was implemented. In the second practice block, a four-up one-down staircase procedure was implemented to establish stimuli that could be used for the “medium” level of difficulty, and the same protocol as the hard staircase was used for each trial. For the “easy” level of difficulty, 20 coherent lines were presented, and no staircase was used to estimate this level. Conditions were included so that the number of coherent lines in the “hard” condition could not exceed 16, and the number of coherent lines in the “medium” difficulty condition could not exceed 18.

Following the staircase estimations, the real experiment began (Fig. 4). In all trials, first, the fixation cross was presented at the center of the screen for 1000ms. Following a 100ms blank, a single group of lines was presented for 150ms at either the center of the screen, or in a peripheral location along the same horizontal meridian, 360 pixels away. To discourage participants from starting with their eyes anywhere other than the fixation cross, 50% of trials presented the lines at the center, 25% of trials presented the patch of lines in a peripheral location on the left, and 25% of trials presented the lines in a peripheral location on the right. After the lines disappeared, participants were required to indicate whether or not there was a group all tilted in the same direction (by pressing Q or P, respectively), and following this, were also required to rate how confident they were in their responses, on a scale from 1 (not at all confident) to 4 (extremely confident).

There were 360 total trials in the main experiment. 180 trials were presented at the center, and 180 trials were presented in peripheral locations (90 left, 90 right). Within each condition (center/periphery), 60 trials were of easy difficulty, 60 medium difficulty, and 60 hard. Catch trials were added at four different trial markers in the experiment (40, 120, 200, and 280). During a catch trial, a letter was displayed at the center of the screen for 1000ms. After the letter disappeared, participants were asked whether an a, b, c, or d was displayed, and were required to input a response on the keyboard. Participants were instructed to take a break for at least 30 seconds after trial 80, 160, 240, and 320.

### Analysis

Each individual participant’s data were analyzed using custom software for Signal Detection Theory (SDT) analysis [44–46]. We used the SDT_MLE_fit.m file to estimate basic Type 1 SDT parameters for sensitivity (*d’a*) and bias (i.e., the criterion*, c’a*) for the aggregated data across all three difficulty levels, and a modified version of the type2_SDT_SEE.m to compute the hit rates and false alarm rates. In Experiment 1, we used the standard SDT measure *d’* because in a discrimination task of that nature, it is unlikely that the equal variance assumption for the two stimulus representations was violated. However, in detection tasks this tends to be an issue. Thus, we used the measure *d’a* to account for potential differences between the variances of the signal and noise distributions [53,54].

## Results

As shown in Figure 5, participants were more sensitive (i.e., exhibited higher *d’a*) in detecting whether a group of lines with congruent orientations was present in the central part of the screen (at fixation), compared to when lines were presented at peripheral locations (t(70) = 7.39, p < 10^−9^). Participants also used different criteria for evaluating whether a coherent patch of lines was present at the center of the screen or the periphery. Specifically, participants were more liberal in detecting coherence in the periphery compared to the center, as shown in the differences in *c’a* (t(70) = 3.89, p < .001). This resulted in a higher number of false alarms (responding “yes” when only random lines were presented) in the periphery compared to center.

**Figure 5.**
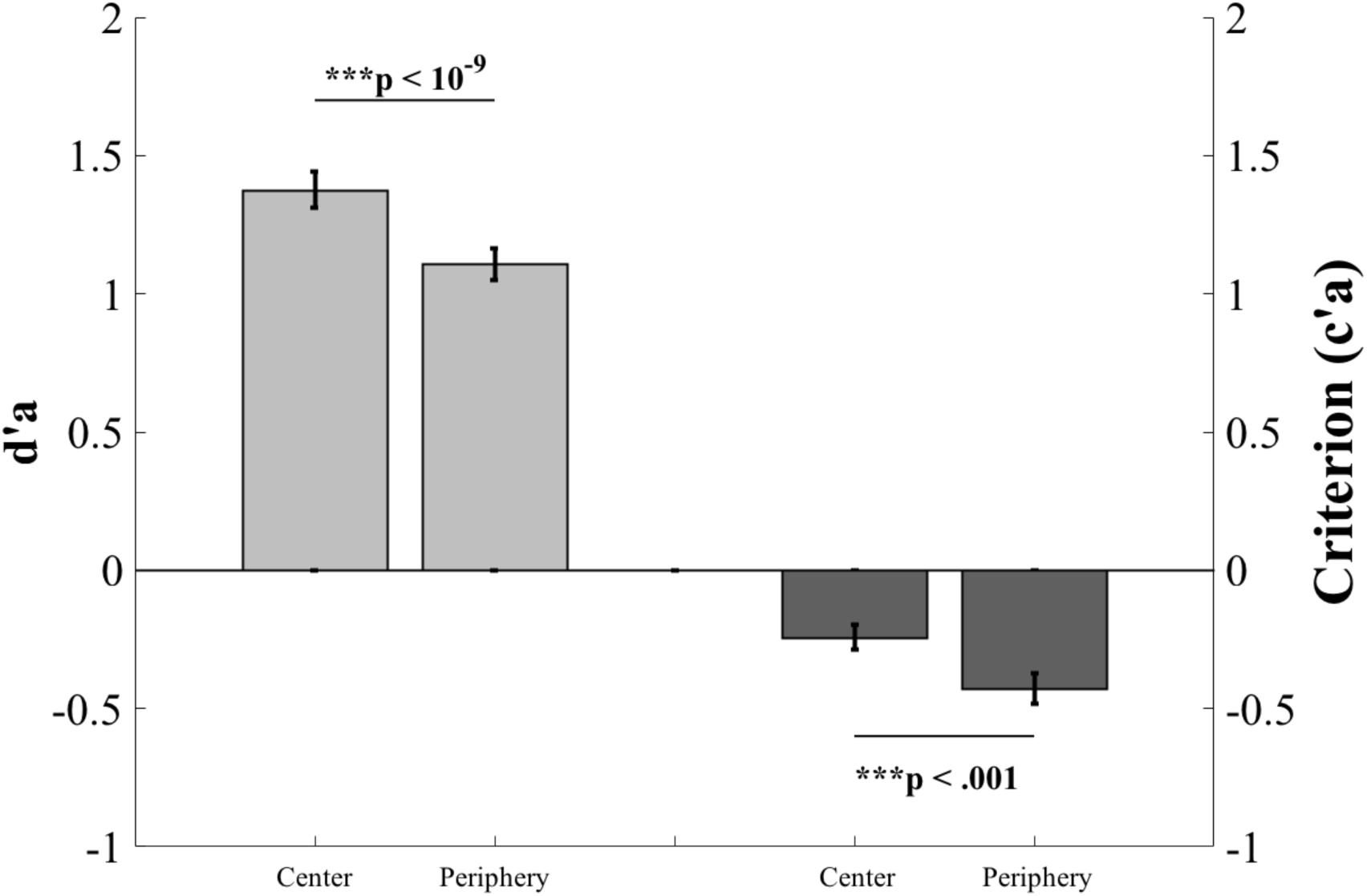
Sensitivity and bias for detecting congruently-oriented groups of lines. Shown here are results from an experiment where participants were asked to detect whether a group of lines with congruent orientations were presented, in either a central or peripheral location. As shown by the light gray panels, using the measure *d’a* (which corrects potential unequal variance in detection tasks), participants were more sensitive in detecting the congruent patch of lines at the central location compared to a peripheral location, and yet they used a more liberal criterion *c’a* in the periphery for indicating that a patch of lines was present. Notice that although sensitivity was not perfectly matched between center and periphery, usually we expect subjects to be relatively conservative for weaker detection, based on the Neyman-Pearson objective [53,55]. Therefore, the results are striking in that it went opposite to that expectation.

Specifically, on trials where lines with only *randomly-sampled* orientations were shown and participants incorrectly reported that a congruent patch was presented (i.e., false alarms), results revealed that participants were much more likely to incorrectly respond when the lines were presented in the periphery, compared to trials where the lines were presented in the center (Fig. 6, t(70) = -6.80, p < 10^−8^). No difference across conditions was found in trials where a coherent patch was presented and participants correctly responded (t(70) = - 0.79, p = 0.43)

**Figure 6.**
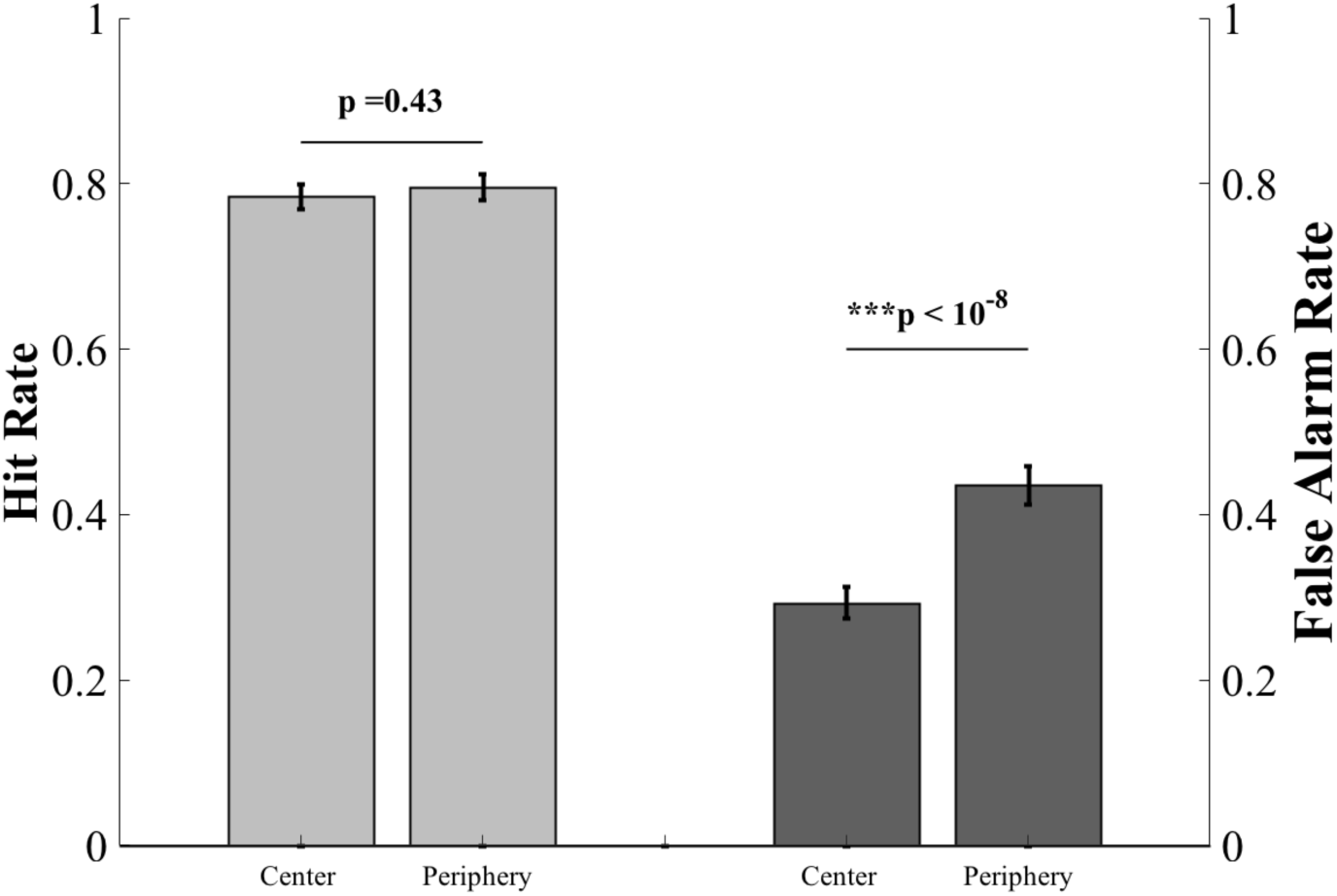
Hit rate and false alarm rate for detecting congruently-oriented groups of lines. In this figure, the light gray bars along the left axis denote the hit rate (i.e., correctly responding that a group of lines with similar orientation was presented) and the dark gray bars along the right axis display the false alarm rate (incorrectly reporting that a group of lines with similar orientation was shown, when only random lines were presented). While the hit rate for detection in this task was quite similar across the two conditions, the false alarm rate was significantly higher in the peripheral condition, compared to the central condition.

These results conceptually replicate previous studies showing that observers use liberal perceptual criteria when making detection-related judgments in the periphery [31,32], and indicate that this liberal detection criterion is used for not only detecting simple stimuli like Gabor patches, but also for more complex stimuli involving summary statistics as well.

## Discussion

We considered how peripheral visual perception may demonstrate *inflation*, whereby subjective judgments in this region of space are marked by two behavioral characteristics: *metacognitive impairments* in how effectively confidence judgments track the correctness of responses in experimental tasks, and *decisional biases* in observers’ tendencies to assume stimuli are more likely to be presented in the periphery than what actually occurs. We conducted two experiments to investigate whether these deficits would emerge in tasks which exploit two well-established phenomena in the visual surround: crowding and summary statistics. In our first experiment using crowded stimuli, observers showed relative deficits in metacognitive measures (e.g the “M-ratio”) [46,47] for crowded compared to single stimuli. This metacognitive deficit was primarily driven by overconfidence in incorrect responses, which is striking given that subjects did not perform the primary discrimination task very well under crowding; the overconfidence is highly unwarranted. In our second experiment using a summary statistical stimulus (groups of oriented lines), observers exhibited liberal detection criteria and high numbers of false alarms, showing that decisional biases extend to more complex stimuli than has been previously shown. Both of these findings provide experimental evidence that, far from perceiving the visual periphery with a high degree of fidelity [3,56,57], our subjective sense of the visual surround is inflated.

Our findings on inflation go beyond what has been shown previously. Previous research has shown that under sensitivity-matched conditions, inflation occurs under inattention and in the periphery [31,32] for simple stimuli, such as Gabor patches. In the present research, sensitivity was not matched, but the stimuli were optimized to exploit the characteristics of peripheral vision, with multiple inputs incorporated simultaneously. These conditions capture the everyday challenges faced by peripheral vision: a deluge of inputs and inherent processing limitations. Under these conditions, the lower sensitivity in crowded/peripheral conditions is expected to result in more conservative detection criteria and lower confidence, but the results strikingly showed the opposite effects. These results demonstrate the prevalence of inflation: it happens also when we did not contrive to fully compensate for the reduced sensitivity in the periphery (by presenting it with stronger stimuli). Inflation is present in various scenarios, including when more complex stimuli are used to challenge the processing bottleneck in peripheral vision. Because crowding and summary statistical judgments happen often in the periphery in everyday life, if inflation is more easily observed in these paradigms than in typical psychophysical experiments involving single targets, the phenomenon may be more prevalent than previously thought [57].

One difference between the present work and previous studies [31,32] is that in our second experiment, our peripheral stimulus incorporated elements of both endogenous and exogenous attention. That is, the peripheral stimulus carried inherent locational uncertainty as it could arise in one of two locations, and this uncertainty likely resulted in not only trial-to-trial differences in the allocation of endogenous attention, but also how exogenous attention may have played a role, too, when the stimulus was presented. That may also explain why the effect of inflation was robust even though sensitivity between center versus periphery was not matched. Future investigations should aim to systematically investigate how exogenous attention and endogenous attention may alter the characteristics of inflation that we observed here.

These findings raise an important question: what may be a mechanistic explanation for inflation? Previously, one proposed account based on SDT is the variance reduction model [31]. According to the model, inattention and peripheral presentation do not drastically alter the perceptual criteria used to make judgments; therefore, the increased variance in internal response in these circumstances causes a greater frequency of occurrences that the response crosses a detection or confidence criterion. Although there are caveats as to whether the criteria are really so inflexibly fixed [58], the model has also been directly tested and highly counter-intuitive predictions have been confirmed [59]. Nevertheless, we acknowledge the simplistic nature of this model. Future work is needed to further elucidate a biologically realistic mechanism.

One potential concern is that based on the results from our first experiment, one could argue that all we observe is a change in confidence, and that the link between confidence and perceptual phenomenology is tenuous. While we acknowledge that confidence is not synonymous with phenomenology *per se*, there are many cases where confidence provides an effective assessment of phenomenology’s presence or absence. For example, in blindsight patients, visual task performance is often spared, but phenomenology is not, and confidence ratings provide an effective means to assess the absence of experience of visual content [60,61]. But we note that even when we ask non-metacognitive questions, as in our second experiment, results indicate that observers think they see more of the periphery than they actually do. It is the joint observation, that peripheral perception leads to both erroneous overconfidence and liberal detection bias, that led us to think these findings may be relevant for subjective phenomenology.

Also, we interpret these findings to reflect inflation, but this is not to say we rule out interpretation based on filling-in as well. Although sensitivity was lower in peripheral detection as well as crowding, such low sensitivity could be the result of filling-in of illusory (i.e.,. non-veridical) content too. Our point here, though, is that over and above potentially filling-in, inflation is likely at play, and its role in accounting for phenomenology in the periphery is at least as important [30].

Finally, why would this sense of inflation have evolved in the visual system? When considering the decisional bias that is present, it becomes important to reflect on what cost functions the visual system may be trying to optimize [62]. A liberal detection bias which causes higher numbers of false alarms may not be “optimal” according to strict signal detection theory. But in a dynamic, changing world which requires fast identification of objects for survival, perhaps a slight overestimation of the presence of objects in the periphery is optimal in the sense that identification of potential threats or rewards can spur exploration and action, to avoid predators and find food and mates. These liberal detection biases may also reflect a larger tendency of perceptual and cognitive systems to make high numbers of false alarms for not only attributes like presence or absence, but also agency in situations where none exists [63]. Overall, these considerations may account for why we subjectively perceive the visual world as relatively uniform despite the poor sensitivity in the periphery.

## Additional Information

### Ethics

All research in Experiment 1 was approved by the Departmental Research Ethics Committee in the Department of Psychology at the University of Hong Kong. Experiment 1 was an undergraduate thesis. All research in Experiment 2 was performed under UCLA IRB #15-001484 and was conducted in accordance with the Declaration of Helsinki.

### Data Accessibility

All code and datasets supporting this article have been uploaded as part of the supplementary materials.

### Authors′ Contributions

M.Y.C. and S.H.C performed the data collection and main analyses for Experiment 1. B.O. conducted the final statistical analysis and created figures for Experiment 1, and performed the data collection, main analyses, and figure creation for Experiment 2, under the guidance of H.L. B.O., M.Y.C., H.L., and S.H.C. wrote the paper.

